# Dietary *Phyllanthus emblica L.* (Indian gooseberry, Amla) improves fecundity and resistance to oxidative stress in a *Drosophila melanogaster* model of early-life malnutrition

**DOI:** 10.1101/2024.12.01.626288

**Authors:** Pallavi Padmaraj, Megha

## Abstract

Amla is a celebrated ethnobotanical fruit whose consumption is associated with several beneficial claims. Many studies have demonstrated Amla’s positive impact on molecular and systemic readouts in diseased conditions. Studies on Amla’s potential as a nutraceutical however are limited. To test if daily dietary supplementation with Amla improves select systemic readouts associated with wellbeing, and alleviate phenotypes associated with early life malnutrition, we deployed the fly animal model system. Benefits were compared between adult flies that were subject to larval starvation (ELS) and controls, and between genders. The most dramatic effect was observed in resistance to oxidative stress: prophylactic feeding of 1% Amla Juice (AJ) increased median survival by ∼75% (control) and 200% (ELS) in males, and ∼167% (control) and ∼150% (ELS) in females, respectively. Interestingly, survival also increased in both genders when AJ was fed only during stress or, before and during stress. For other assays, impacts were seen only in ELS flies: AJ feeding decreased lifespan in female ELS flies, increased egg numbers by ∼25% and improved survival upon *P. entomophila* infection by ∼80%. Together, these suggest that prophylactic AJ dietary intake has selective biological effects and in the context of malnutrition, it can be explored further as a nutraceutical.

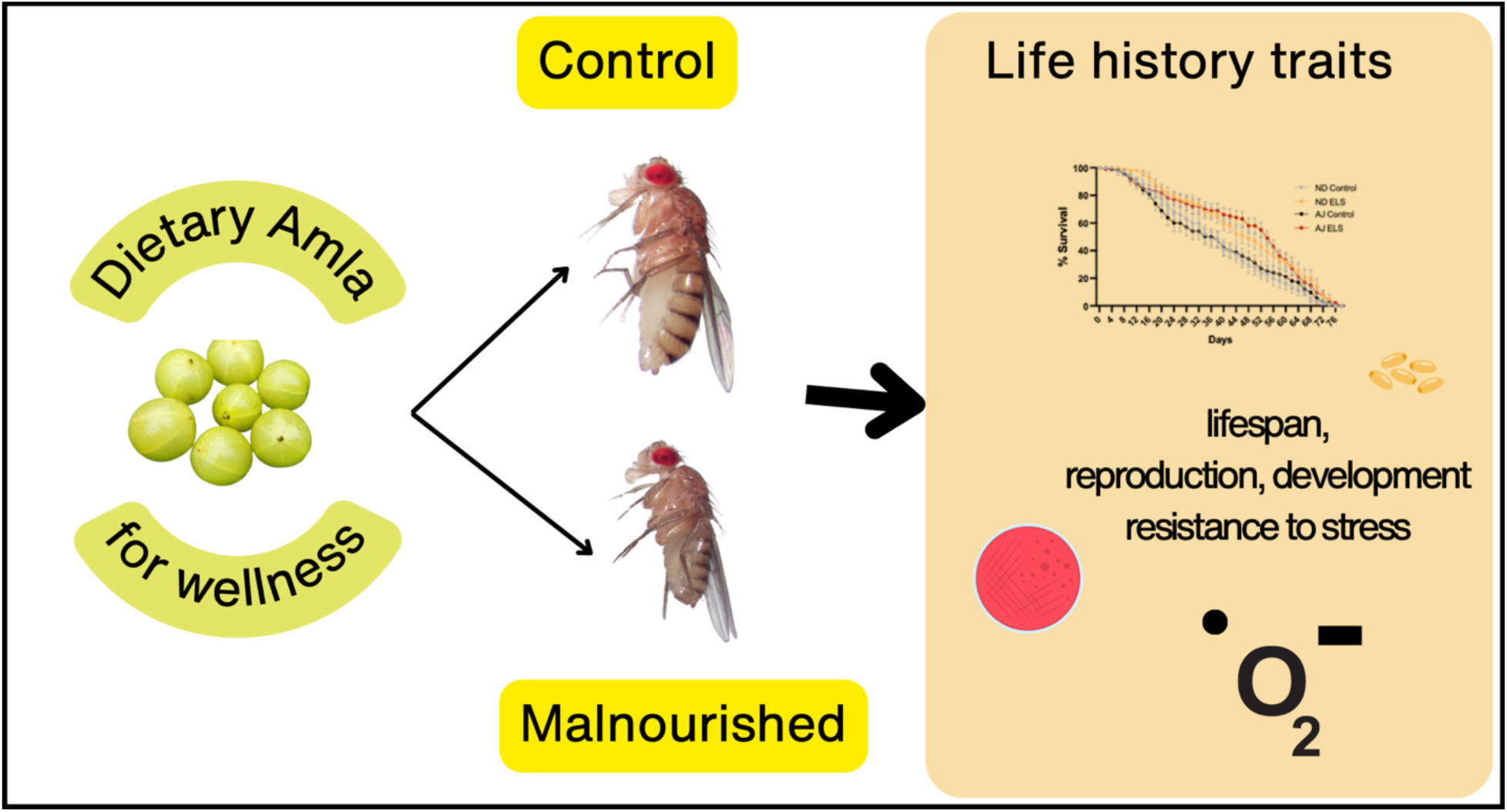

## 1. Introduction

Dietary supplements are a popular means of achieving optimal nutrition to sustain good health. Consumer preference for natural ingredients has led to the addition of food ingredients associated with benefits for wellbeing purportedly supported by traditional medical claims. Yet, in a majority of cases, benefits testing for individual ingredients has primarily focused on outcomes in diseased contexts, not wellness. The fruit of *Phyllanthus emblica L (Indian gooseberry)* [Sanskrit: Amla; Synonym: *Emblica officinalis Gaertn*] is one such extensively studied and consumed medicinal food. In codified and non-codified traditional Indian medical systems, Amla is believed to be anti-inflammatory, anti-dysentery, hepatoprotective, neuroprotective, antidiabetic, microbicidal and immunomodulatory (Ahmad et al., 2021; Ma et al., 2024). The fruit by itself, and as a part of several formulations is prescribed in traditional medicine for diverse health benefits, from general well-being to the management of conditions like infertility, diabetes, and iron-deficiency anaemia (Patel & Goyal, 2011). Interestingly, although the fruit is more popularly consumed, nearly all parts of the plant such as leaves, seed, bark, and flowers, are utilized in traditional remedies (Gul et al., 2022; Masihuddin et al., 2019; Mirunalini, & Krishnaveni, 2010; Prananda et al., 2023).

Amla is both the name of the fruit and whole tree bearing the fruit. It belongs to the family *Phyllanthaceae*. The weight of the fruit varies from 25.94 to 33.90g and the length varies from 3.07 to 3.82cm ((Avinash et al., 2024); Supplementary Fig. 1). Fresh fruits are light green in colour, turning to light brown upon ripening. It is native to Southeast Asia and extensively cultivated in India, Nepal, Sri Lanka, Pakistan, and Bangladesh (Avinash et al., 2024). The fruit is an abundant source of bioactive chemical groups such as tannins, flavonoids, alkaloids, organic acids and polyphenols, which have been shown to act as antioxidants and free-radical scavengers (Ahmad et al., 2021; Khopde et al., 2001; Poltanov et al., 2009).

Noteworthy among these are hydrolysable tannins such as Emblicanin A and B, gallic acid, and ellagic acid, which have demonstrated significant biological activity (Gul et al., 2022; Masihuddin et al., 2019; Mirunalini, & Krishnaveni, 2010; Patel & Goyal, 2011). Another notable constituent of the fruit is vitamin C, a potent antioxidant, which has been reported to be present in fresh fruit in concentrations ranging from 400mg (Scartezzini et al., 2006)-252mg (Longvah et al., 2017) per 100g of fruit. Additionally, Amla is enriched with essential nutrients including phosphorus, iron, sodium, vitamin A, fibre and magnesium, further bolstering its therapeutic value (Longvah et al., 2017).

A meta-analysis of nine randomised control trials (RCTs) where Amla was tested as a dietary supplement in pill form concluded that its consumption is associated with improved blood lipid biomarkers; however clinical heterogeneity and small sample size were noted to preclude strong statements on the efficacy (Brown et al., 2023). For instance, two RCTs using a patented pill containing 500mg of Amla powder (CAPROS, USA) are reported in diseased contexts. In one where n =15, an improvement on cardiovascular risk factors such as collagen induced platelet aggregation and the inflammatory marker, hsCRP, was reported in obese persons (Khanna et al., 2015). The other, where n = 59, reported improved endothelial function, increased levels of serum markers of anti-oxidative molecules (nitric oxide, glutathione) and minor improvements in lipid biochemistry in persons with metabolic syndrome (Usharani et al., 2019). In healthy volunteers consuming Amla, a small randomised (n=15) trial in Japan showed improved vascular function and haematological parameters, as well as lipid profile (Kapoor et al., 2019). Altogether, these studies indicate that benefits of Amla on human biological readouts requires more rigorous trials, but one aspect that stands out is that there is no toxicity or adverse effects associated with Amla consumption.

Several in vitro and in vivo studies have demonstrated that Amla contains bioactive molecules that interact with diverse signalling pathways and enzyme systems involved in mitigating disease outcomes. In chemical-induced rodent cancer models, an aqueous as well as polyphenol extract reduced tumour progress and improved survival outcomes (Kumar et al., 2022). Hepatoprotective benefits have been demonstrated in rats with carbon-tetrachloride- (CCl_4_-) induced liver fibrosis (Yin et al., 2021), alcohol-induced liver damage (Damodara Reddy et al., 2010) as well as in the HepG2 cell line (Thilakchand et al., 2013). Anti-inflammatory activity of a hydroalcoholic extract of Amla reportedly reduced carrageenan-induced (and other phlogistic agents induced) paw oedema in rats and, at 700mg/kg provided as much pain relief as over-the-counter NSAIDs (Golechha et al., 2014). Significant analgesic effects were also reported in plantar incision and spared nerve injury rat models of pain where Amla supplementation correlated with increased mechanical withdrawal thresholds and reduced pain-related vocalizations (Lim et al., 2016). At the cellular level, a fruit extract powder of Amla was shown to scavenge reactive oxygen species produced in RAW246.7 macrophages upon lipopolysaccharide stimulation, as well as suppressing NF-κB, COX-2, and iNOS expression (Wang et al., 2019). Cumulatively, these studies point to constituents of Amla modulating inflammatory cytokines such as IL-6, NF-kB, enzymes such as superoxide dismutase, and signalling pathways involved in oxidative stress as well as the cell cycle (Kumar et al., 2022; Thilakchand et al., 2013). A recent study using an in silico network pharmacology of Amla illustrates diverse biomolecular targets that are spread across various signalling pathways, underscoring the poly-pharmacological nature of the Amla fruit (Agarwal et al., 2023).

A common narrative in the vast majority of published studies on the utility of Amla is a focus on its therapeutic value in the context of diseases ranging from cancer, obesity, hepatic disorders, to heart diseases. However, both in classical Ayurvedic texts, one of India’s medical knowledge systems, as well as contemporary culture, Amla is celebrated as a dietary ingredient that promotes wellness (Ruhela et al., 2023). This aspect of Amla, as a prophylactic that can be consumed daily as a food rather than medicine, is less well-researched. For whole animal testing, where Amla can be introduced into the diet without compromising its format as a freshly expressed juice, we turned to *Drosophila melanogaster* (fruit/vinegar fly). Flies are an attractive model system to test dietary interventions due to their short lifespan, genetic similarities with humans in fundamental metabolic pathways, and cost-effective maintenance (Eickelberg et al., 2022; Verheyen, 2022). Its rapid life cycle enables efficient assessment of dietary impacts across generations, making it ideal for studying long-term effects on fecundity, aging, and longevity (Bazzell et al., 2013; Dwivedi & Lakhotia, 2016; Klepsatel et al., 2020; Savola et al., 2021). A primary advantage of the fly system is that fly food is cooked similar to human food as a porridge of corn flour, yeast, and sugar, and that dietary interventions can be easily introduced to this mix (Eickelberg et al., 2022). The fruit fly’s versatility in diet manipulation facilitates controlled experiments on calorie intake, nutrient composition, and supplementation, to help elucidate how dietary components can influence a variety of biological processes from development to disease susceptibility (Bazzell et al., 2013; Verheyen, 2022; Yi et al., 2021).

Amla as a potential dietary intervention to address malnutrition, a systemic issue that can result from a lack of quality nutrients or diseases such as cancer, remains unexplored. Globally, more than 149 million children are estimated to be either stunted or wasted, with maternal and early childhood malnutrition being a major cause (Kinyoki et al., 2020). Childhood malnutrition is tackled through a slew of policy and material interventions, of which diet is an integral component. To provide an impetus to this important public health issue, we wished to understand how dietary intervention with Amla can impact adult phenotypes in a fly model of early-life malnutrition developed in our lab (Patil et al., 2022). By manipulating yeast concentrations during larval development, studies have reported an impact on adult phenotypes such as fecundity, longevity, resistance to infection, and lipid metabolism (Collins et al., 2023; Klepsatel et al., 2020; Rehman & Varghese, 2021; Savola et al., 2022; Stefana et al., 2017). In this work, we report similar phenotypes, and test how dietary supplementation with Amla can ameliorate these phenotypes.

## 2. Materials and Methods

### 2.1 Preparation of freshly expressed amla juice (AJ)

Fresh Amla fruits (Indian gooseberry/ *Phyllanthus embilica L;* 35 ± 3 mm diameter, 27 ± 2 g) were either purchased from the market or when in season, obtained from the TDU campus medicinal garden. The fruits were washed thoroughly and surface sterilized using 70% ethanol. Deseeded, chopped fruits were then pulped using Dr. Mills grinder chopper (DM-7412M) and the juice was strained through a clean muslin cloth. Amla Juice (AJ) obtained from 3-4 fruits was aliquoted and stored at −80°C until use. 5% of fresh AJ hence prepared had a scavenging activity of 160%, suggesting an equivalence to 0.01mmoles Ascorbic acid, which appeared not to change upon storage for 3 months at -80°C (Supplementary Table 1). Yet, we found that samples more than a month old gave variable results in the fly experiments, therefore for all experiments, AJ used is not more than 3-4 weeks old.

### 2.2 Fly husbandry

*Canton-S* strains of *Drosophila melanogaster* were used. The flies were reared in a controlled environment with a 12 h:12 h light-dark cycle at 25°C. Flies were raised on media (Normal diet; ND), 1000mL of which consists of 80g Corn meal (locally sourced), 40g D-Glucose (Qualigens-Q15405), 20g Sucrose (Qualigens-Q15925), 50g Yeast extract (HIMEDIA-RM027), 8g Agar (Qualigen-Q21185), 4mL propionic acid (Qualigen-Q26955), 7mL Benzoic acid (Qualigens-211665), and 6mL of 85% ortho-phosphoric acid (Qualigen-Q29245). Amla juice (AJ) media was prepared by addition of undiluted AJ upto 1% (v/v) to the cooked ND as it was cooling. Adult flies were fed AJ right from the day that they eclosed.

### 2.3 Generation of Early Life Starved (ELS) flies

The ELS flies were generated as detailed previously (Patil et al., 2022). In brief, flies were allowed to lay eggs for 4-6 hours. Subsequently, L3 larvae were collected after 90 hours, cleaned and transferred to either ND media or 100 mM Sucrose. After eclosion, the adult flies were transferred to AJ media or ND media, with 25-30 adults per vial.

### 2.4 Longevity assay

Newly eclosed adult flies were collected and transferred into vials with AJ or ND media. Sexes were cohoused during the experiment. Flies were transferred to fresh media every second day and the number of dead flies was counted every day.

### 2.5 Fecundity assay

5-days old females (5-7) and males (3-4), matured on different diet regimens were taken per vial. The total number of eggs laid every 24 hours for 3 days were counted and averaged by day, and female number.

### 2.6 Development assay

Flies were allowed to lay eggs for approx. 4 hrs to keep egg numbers at approximately 30 per vial. The total number of eggs, pupae (every 6 hours after the first pupae were spotted), and as time went, adult numbers were counted, till at least 14 days post egg laying.

### 2.7 Paraquat-induced oxidative stress assay

7-day-old flies fed on respective diet regimes were sorted by sex and then exposed to 5mM paraquat (PQ, Sigma, Catalog no 36541) dissolved in 5% sucrose and in controls, to only 5% sucrose. For exposure to Amla during stress, 1% AJ (v/v) was mixed along with the paraquat solution. The number of survivors was counted every 12 hrs, for upto 10 days. For qRT-PCR experiments, 5-day-old flies fed on respective media were given PQ treatment for 48 hours, after which their RNA was isolated.

### 2.8 RNA isolation, cDNA synthesis and quantitative Real Time-PCR (qRT-PCR)

RNA extraction was performed on whole flies (3 samples per group) using Trizol-chloroform (RNAiso Plus-TaKaRa 99108, Chloroform 99.5% SRL-84155). cDNA was synthesised using the Takara PrimeScript™ RT Reagent Kit (Perfect Real Time, RR037A). qRT-PCR was performed using the following gene-specific primers shown below. Relative changes in gene expression were calculated using the ΔΔC_t_ method and expressed as fold changes relative to *rp49,* a housekeeping gene.

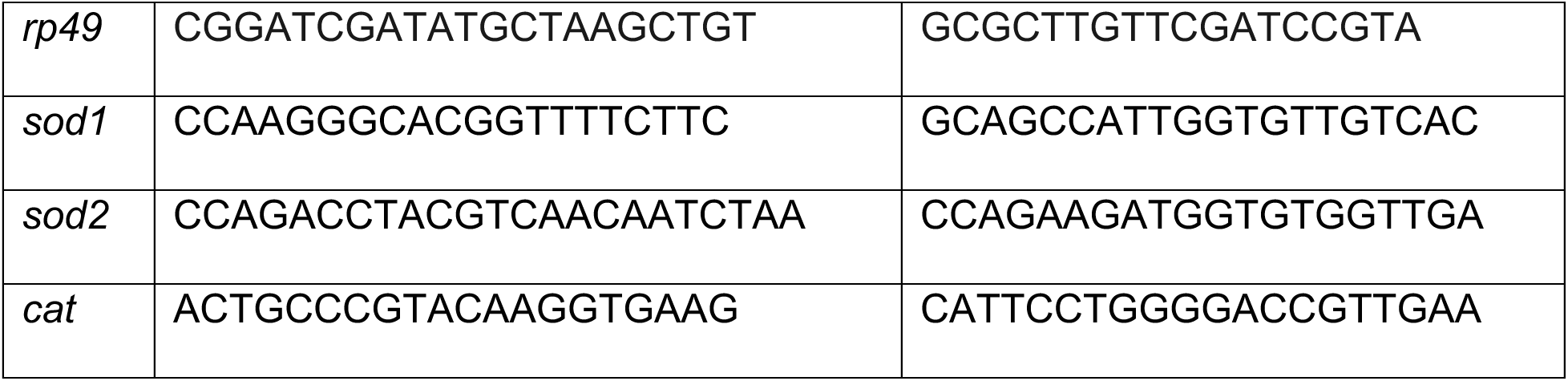

### 2.9 Infection with Pseudomonas entomophila

For enteric infections, the protocol described by Buchon et al., 2010 (Buchon et al., 2010) was adapted. 7-day-old flies were starved for 2 hours, followed by a 24hr incubation with a filter disc containing an overnight culture of *Pseudomonas entomophila* at OD_600_ = 150 in 5% sucrose. *Pseudomonas entomophila* was a kind gift from Dr. N. G. Prasad, IISER Mohali. When AJ was provided during infection, it was mixed at 1% (v/v), along with the bacterial solution. Flies were returned to ND media vials 24 hrs post-infection and were maintained at 30°C thereafter. Survival was monitored till day 5 post-infection.

### 2.10 Statistical Analysis

Sample size and numbers are indicated in the figure legends. Plots have been graphed and statistical comparisons performed using GraphPad Prism 8.4.2 software. Statistical analysis for the survivorship curves in the longevity assay and the PQ-induced oxidative stress assay was performed using Kaplan-Meier analysis with log-rank post-test. The median survival of flies in the longevity assay and PQ-induced oxidative stress was analysed using Mann Whittney U test. Student’s t-test was performed to compare the mean between groups in the fecundity and viability assays. One-way ANOVA was performed to analyse the percentage survival upon enteric infection.

## 3. Results

### 3.1 Impact of dietary supplementation with Amla Juice (AJ) on select life-history traits

Phenotypes such as lifespan, reproduction, and development are standard measurable outcomes used to understand how external stimuli such as diet, can effect organismal health. Accordingly, we first tested how chronic dietary supplementation of AJ regulates longevity, egg numbers, egg viability and pupal developmental time.

As reported previously (Koliada et al., 2020), mated males had longer median survival times than mated females and here we find this persists regardless of larval stress (Fig. 1A and 1B, Supplementary Table 1). In controls, feeding AJ had neither a negative nor a positive impact on lifespan. This contrasts with previous reports that Amla feeding in flies results in lifespan extension <u>(Pathak et al., 2011; Rawal et al., 2014)</u>. These studies however do not detail the dosage or method of Amla processing, as well mating status of the flies, thus limiting the ability to make comparisons. AJ feeding did not increase the longevity of male control and ELS flies. In contrast, AJ feeding mildly decreased longevity of ELS female flies, with median survival falling from 18±1.64 to 16±0.63 days (p < 0.0001), bringing it in line with control females fed AJ, at 16±0.88 days. To summarise, an impact of AJ feeding on lifespan was observed only in mated ELS flies and is sex-dependent.

**Figure 1:**
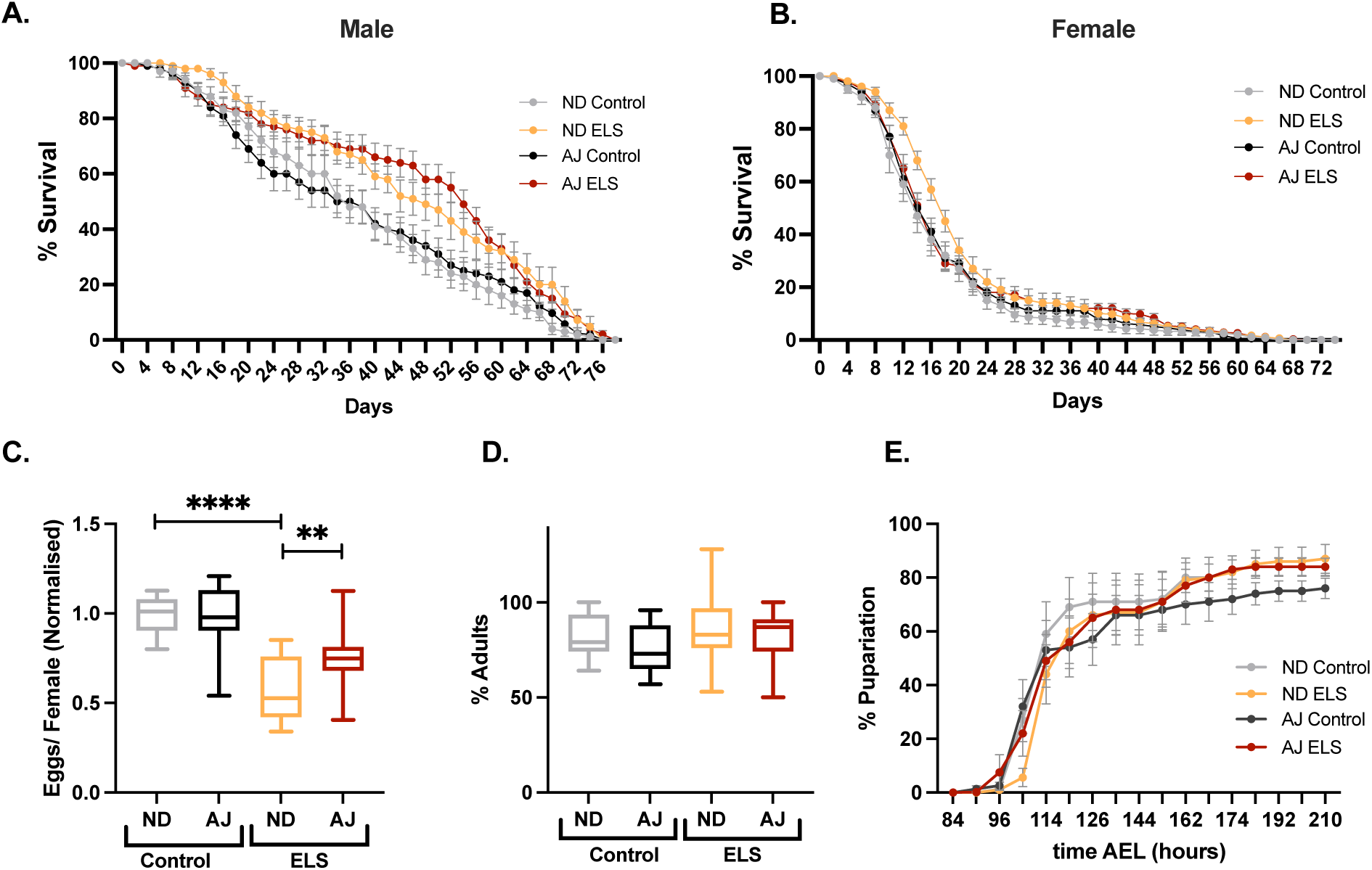
Selected life-history traits of flies on normal diet (ND) or ND supplemented with 1% Amla Juice (AJ). Larvae collected ∼ 94 hrs after egg-laying were placed in 100mM Sucrose [Early life Starved (ELS)] or ND [Control] and adults emerging from respective media were tested in these assays. **A.** Males: Lifespan of control and ELS flies on ND or AJ media. n = Control ≥ 176, ELS ≥ 134, p values: ND control vs ND ELS, p = 0.0001; ND ELS vs AJ ELS, p = 0.65 **B.** Females: Lifespan of control and ELS flies. n: Control ≥ 291, ELS ≥ 269. p values = ND control vs ND ELS, p = 0.0016, ND Control vs AJ Control p = 0.72, ND ELS vs AJ ELS = 0.0337. Error bars are ± SEM. Statistical analysis was performed using Kaplan-Meier analysis with log-rank post-test. **C.** Average number of eggs deposited per female over 24 hours normalized to ND control values. No of vials =15 **D.** Egg viability as measured by the percentage of adult flies developing from eggs laid on ND or AJ media of either control or ELS background. No of vials = 12. **E.** Developmental time as measured by the time taken for the eggs laid on ND or AJ media by Control or ELS females to develop into pupae. No. of vials = 12. Error bars are ± SEM. Differences between bars are statistically insignificant. ** p ≤ 0.01 and **** p ≤ 0.0001 calculated using Student’s t-test.

It is well known that both nutrient quality and quantity influence reproductive potential in flies. For e.g., reducing yeast concentrations, a critical source of essential amino acids, vitamins, and lipids in the media, significantly reduces egg numbers (C. Ma et al., 2022; Mirth et al., 2019; Simmons & Bradley, 1997; Zanco et al., 2023). If AJ supplementation is beneficial, we hypothesized that egg numbers will increase.

However, this increase was observed only in ELS flies where the average normalised egg numbers went up from 0.56±0.04 to 0.75±0.04, an increase of approximately 25% (Fig. 1C). Interestingly, an increase in reproductive fitness (Fig. 1C) and a decrease in female longevity (Fig. 1B) is a widely recognised reproduction-lifespan trade-off in the field of evolution, and reported across species (Maklakov & Chapman, 2019). An increase in insulin signalling is known to drive this trend in flies (Armstrong, 2020; Grandison et al., 2009; Zanco et al., 2021), which raises the possibility that AJ stimulates the insulin signalling pathway. However, the absence of any impact of AJ on lifespan or egg numbers in control flies argues against this explanation. Instead, perhaps constituents of AJ likely work by inhibiting pathways that repress egg production, rather than stimulating pathways that increase egg production. One constituent of Amla, which has recently been shown to increase egg numbers in flies is vitamin C (Shrestha et al., 2023). In this study (Shrestha et al., 2023), the inclusion of vitamin C at a concentration of 50mM increased egg numbers by ∼4-fold. However, at the concentration (1% v/v) deployed in our assays, vitamin C levels are in the order of 0.1-0.5μM. Also, AJ used in our assays is subject to heat from cooking as well as exposure to air during media storage, and incubation: actions which will reduce vitamin C levels to even lower amounts. Hence, we hypothesize that components of Amla juice, other than vitamin C are regulating egg-numbers.

Next, we measured development-related outcomes: egg to adult viability (Fig. 1D) and pupal developmental time (Fig.1E). Environmental stresses such as poor diet quality, increased temperature and reduced oxygen reduce egg viability, and increase developmental time. It was found that AJ supplementation did not impact either egg-to-adult viability (Fig. 1D), or egg-to-pupae developmental time (Fig. 1E), both in control as well as ELS flies. This suggests that at the concentrations deployed, AJ is neither beneficial nor harmful to development.

### 3.2 Feeding Amla juice significantly improves resistance to oxidative stress

Amla as a whole, its extracts as well as its individual constituents such as vitamin C, gallic acid and quercetin, have been shown to scavenge free radicals and possess antioxidant activity in vitro (Poltanov et al., 2009; Ruhela et al., 2023; Scartezzini et al., 2006; Somasekhar et al., 2016). In rodent studies, both aqueous and alcoholic extracts have been shown to protect against environmental and drug-induced oxidative stress (Anilakumar et al., 2004; Damodara Reddy et al., 2010; Sharma et al., 2009), predominantly in the context of cancer (Singai et al., 2024; Sultana et al., 2008). In flies, there is limited information on the action of whole amla feeding on oxidative stress; however, the positive effect of the phytoconstituents of Amla such as gallic acid, ellagic acid and vitamin C have been reported (Bonilla et al., 2006; Chahal et al., 2020; Packirisamy et al., 2018; Park et al., 2012; Prananda et al., 2023). As ethnobotanical and cultural uses suggest that whole Amla fruit intake can have prophylactic benefits, we deployed a paraquat (PQ) induced oxidative stress assay, under different feeding regimens (Fig. 2A) to test this claim. This assay measures the survival of flies upon continuous exposure to PQ, a herbicide that generates free radicals that overwhelms the fly system, resulting eventually in fly death (Hosamani & Muralidhara, 2013; Rzezniczak et al., 2011). Longer the survival under oxidative stress, better is the anti-oxidant potency of the dietary intervention. Three feeding paradigms with AJ were tested: 1) Pre-stress: flies were fed an AJ-supplemented diet for 5 days before exposure to PQ, a prophylactic feeding condition; 2) During stress: flies were fed AJ only upon exposure to PQ and 3) Pre & during stress: a combination of (1) and (2) i.e., flies were continuously exposed AJ (Fig. 2A). The control dietary regimen was without AJ.

**Figure 2:**
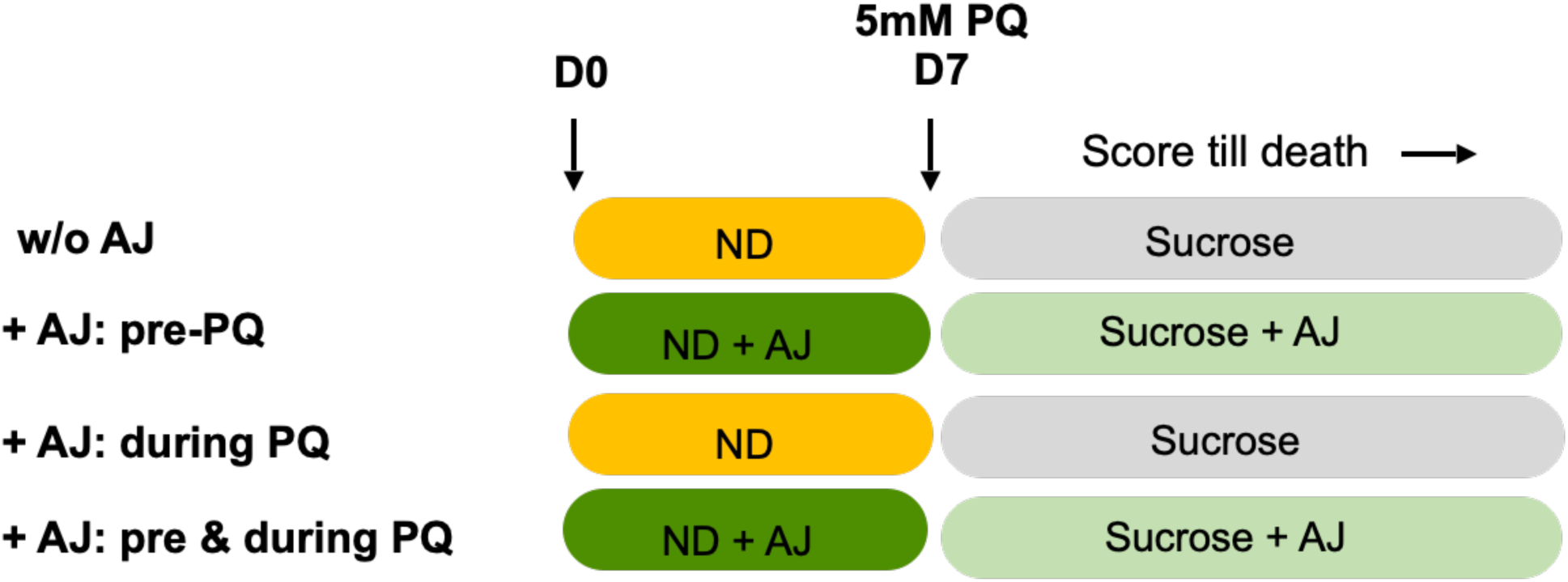
A. Schematic of the four feeding regimens tested in the study. w/o = without; AJ = Amla Juice; PQ = Paraquat; D= Day #; ND= Normal diet. 5mM Paraquat is dissolved in 5% Sucrose. Flies were scored for survival for up to 14 days post exposure to PQ.

We observed that an AJ-supplemented diet drastically improved resistance to oxidative stress, regardless of gender, developmental nutritional stress (control vs ELS) or feeding regimen (Fig. 3A-F, 4A-F; Summarised in Table 1). The percentage increase in survival ranged from 75±38% to 525±20% (Table 1). As previously reported, sexual dimorphism was observed in survival to PQ exposure, with mated females being more sensitive than mated males (Lashmanova et al., 2015; Minois et al., 2012; Pomatto et al., 2017). Interestingly, on diets without AJ, control males performed better than ELS males, while ELS females performed better than control females, such that in ELS flies, males and females had similar median survival of 48 hours (Table 1). Furthermore, dietary AJ supplementation was far more effective on ELS males and control females. In regards to feeding regimen, an AJ-supplemented diet before PQ exposure was sufficient to increase survival resilience (Fig. 3A, D and Fig. 4A, D) in all groups. This supports the notion that AJ has prophylactic potential. When exposed to AJ only during PQ exposure, flies displayed enhanced survival over the pre-stress feeding regimen, except for ELS females (Fig. 3B, E and Fig. 4B, E). The presence of AJ, pre- and during stress resulted in an additive impact on survival for all groups, with this feeding regimen showing the maximum increase in survivorship (Fig. 3F and Fig. 4C, F, Table 1), except for control males (Fig. 3C).

**Figure 3.**
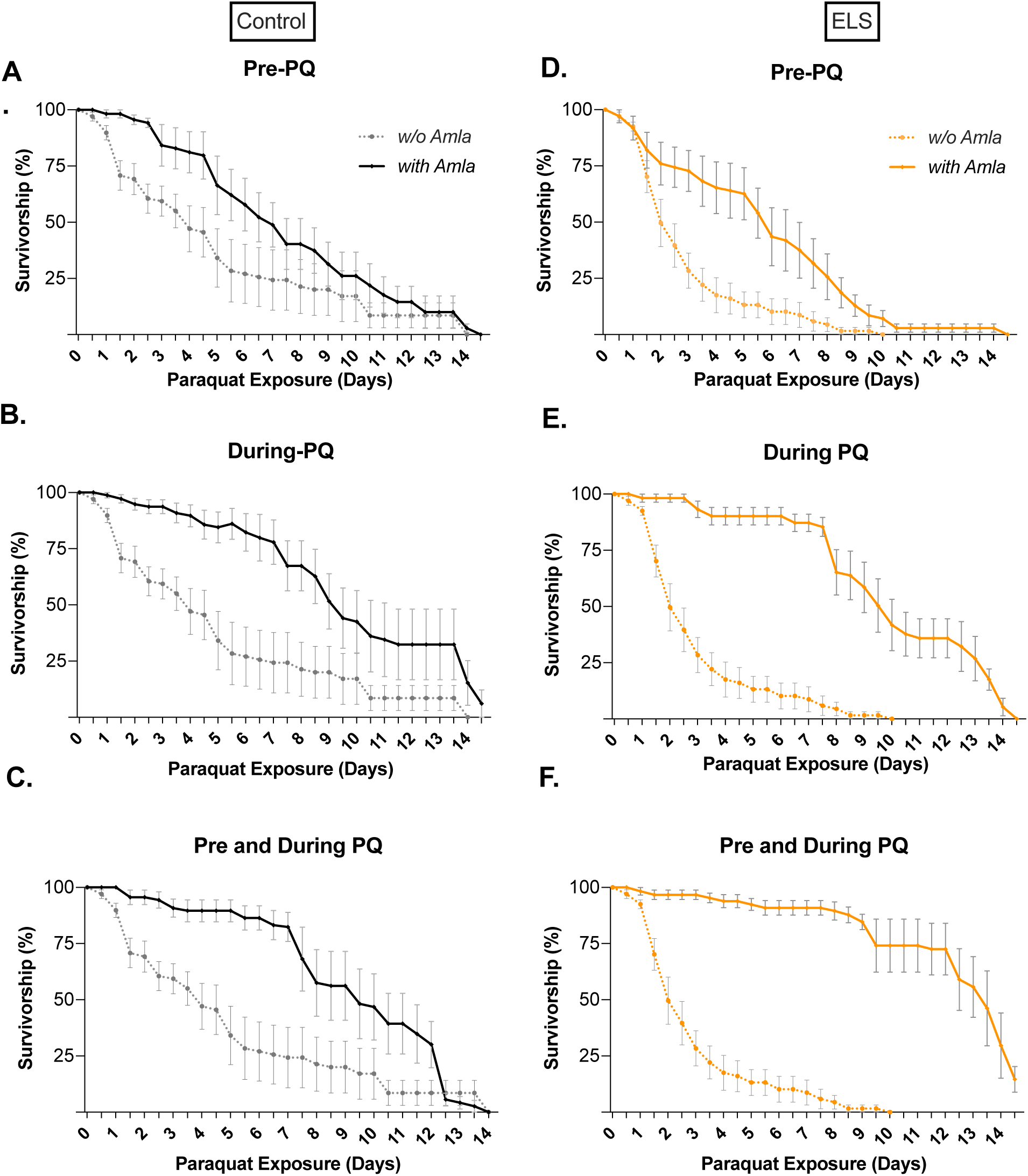
Amla juice improves the survivability upon PQ-induced oxidative stress in males. PQ-induced oxidative stress in control (black) and ELS (orange)-n: Control ≥ 64, ELS ≥ 54). Statistical analysis was performed using Kaplan-Meier analysis with log-rank post-test, Error bars are ± SEM, p value< 0.001. Median survival is tabulated in Figure 2.

**Figure 4.**
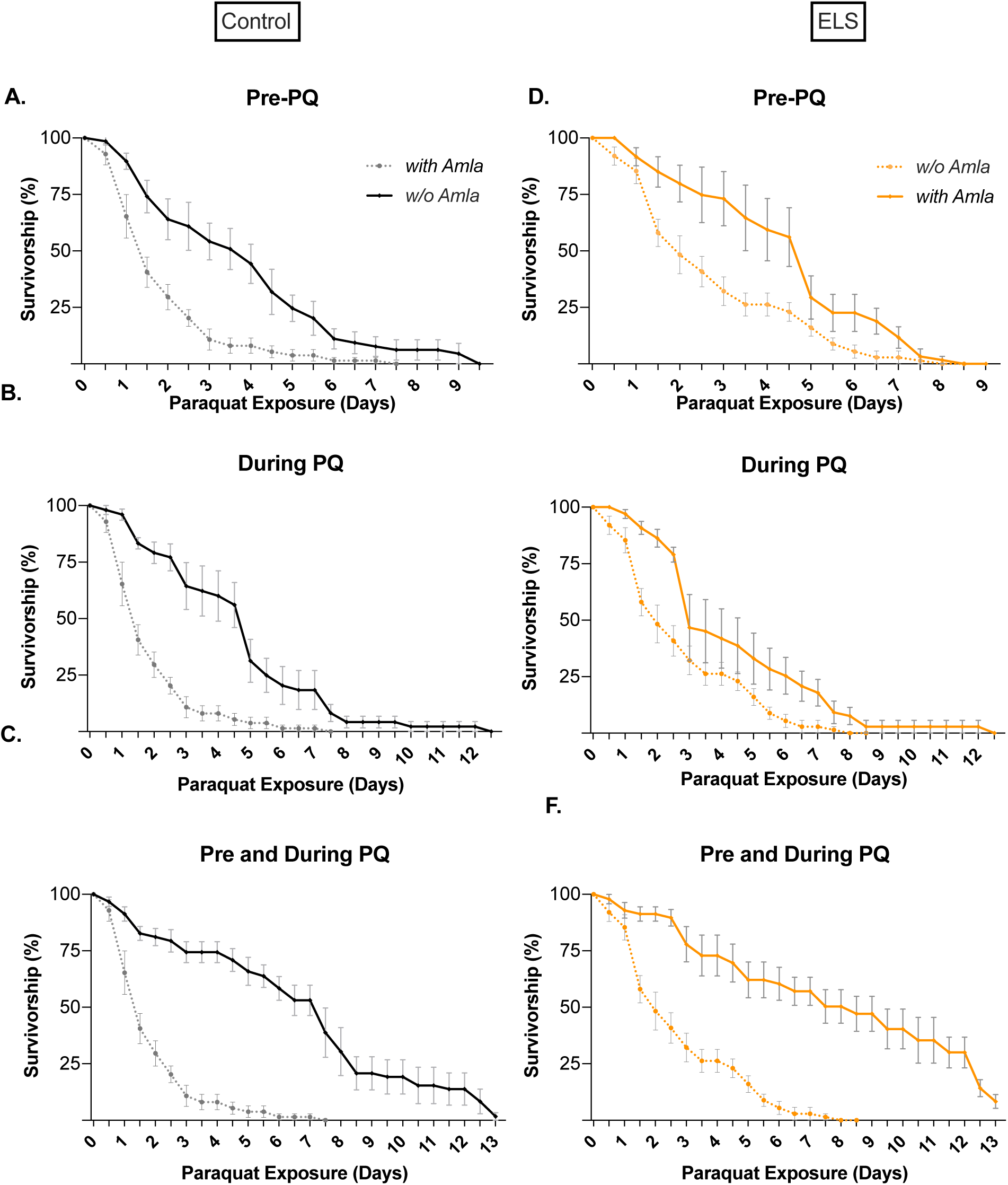
Amla juice improves the survivability upon PQ-induced oxidative stress in females. PQ-induced oxidative stress in control (black) and ELS (orange)-n: Control ≥ 48, ELS ≥ 53). Statistical analysis was performed using Kaplan-Meier analysis with log-rank post-test, Error bars are ± SEM, p value< 0.001. Median survival is tabulated in Figure 2.

**Table 1:** Median survival times for flies exposed to 5mM Paraquat under different feeding regimens (Fig. 2). This table summarises the data presented in Fig. 3 and Fig. 4. Across gender and genotype, differences in median survival were statistically different for all except control males indicated with * and ELS males vs females on a diet without Amla Juice (w/o AJ), indicated by #, when tested by Mann Whitney U test and Kaplan-Meier analysis with log-rank post-test.

| Median Survival Upon Paraquat Exposure (hrs ± SEM) | | | | | | | | |
| --- | --- | --- | --- | --- | --- | --- | --- | --- |
|  | Males |  |  |  | Females |  |  |  |
| Diet | Con | % increase over w/o AJ | ELS | % increase over w/o AJ | Con | % increase over w/o AJ | ELS | % increase over w/o AJ |
| w/o AJ | 96±28 | | 48±6<sup>#</sup> | | 36 ± 6 | | 48±7<sup>#</sup> | |
| + AJ: pre-PQ | 168±26 | 75±38 | 144±25 | 200±26 | 96 ± 12 | 167±13 | 120±13 | 150±15 |
| + AJ: During PQ | 216±32* | 125±43 | 240±20 | 400±21 | 120±12 | 233±13 | 72±12 | 50±14 |
| + AJ: Pre & During PQ | 228±21* | 138±35 | 300±19 | 525±20 | 180±15 | 400±16 | 180±34 | 275±35 |

PQ exerts its toxicity by creating reactive oxygen species (ROS) that overwhelm the cellular redox cycling machinery in hallmarks including increased malondialdehyde, an end product of lipid peroxidation (Hosamani & Muralidhara, 2013; See et al., 2022), reduced dopamine (Fahim et al., 2013; Kang et al., 2009) and increased nitric oxide radicals (Blanco-Ayala et al., 2014; See et al., 2022) as well as an up-regulation of cellular death pathways that eventually lead to cellular death and consequently, animal death (Blanco-Ayala et al., 2014; Chen et al., 2021; See et al., 2022). Oxidative stress at the cellular level is managed by systems that convert the harmful ROS to water: these include, but are not limited to, enzymes such as superoxide dismutase, catalase and redox cycling via glutathione. The ability of components in AJ extracts to scavenge ROS have been established by several studies (Dwivedi & Lakhotia, 2016; Khopde et al., 2001; Poltanov et al., 2009; Scartezzini et al., 2006; Usharani et al., 2019) (we also observed this in our AJ samples; Supplementary Fig. 1), hence we explored if AJ could also exert its effect via biological regulation of superoxide dismutase (*sod1, sod2*) and catalase (*cat*). As previously reported (Park et al., 2012), exposure to PQ reduces the gene expression of cytosolic SOD (*sod1*), mitochondrial SOD (*sod2*) as well as catalase (*cat*) (Fig. 5). Unlike quercetin or *Sanguisorba officinalis* (Park et al., 2012) AJ feeding did not impact the expression of *sod1, sod2,* or *cat*, in either control or ELS flies (Fig. 5). Hence, the prophylactic benefits of Amla intake are likely through other biological mechanisms which require further investigation.

**Figure 5:**
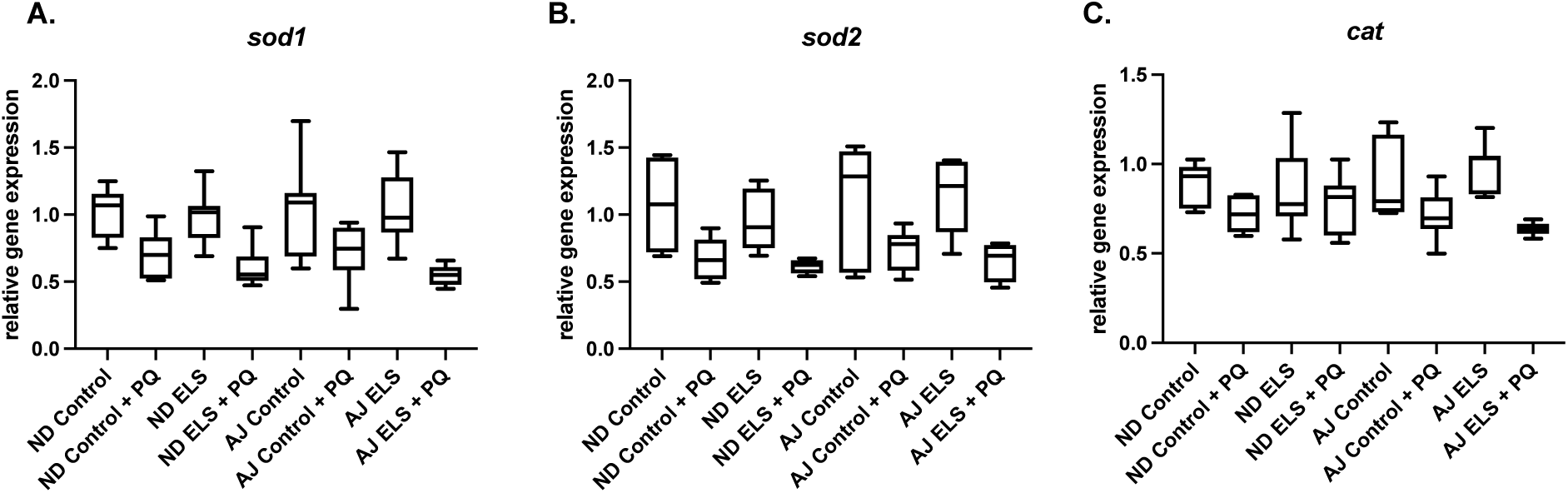
The impact of Amla feeding on gene expression of select enzymes that manage cellular oxidative stress. Relative gene expression of A) Superoxide dismutase 1: sod1 B) Superoxide dismutase 2: sod2 c) Catalase: cat measured before and after 48hrs of PQ exposure (n=7).

### 3.3 Impact of Amla juice supplementation upon enteric infection

The observation that prophylactic feeding with AJ improves resistance to oxidative stress made us wonder whether this positive effect extends to other types of stressors. Dealing with bacterial infection is one such stress; also, it was chosen as there are a number of ethnobotanical and cultural references to Amla’s ability to increase immunity (Jantan et al., 2019; Mirunalini, & Krishnaveni, 2010; Ruhela et al., 2023). Flies were subjected to enteric infection with *Pseudomonas entomophila,* a gram-negative bacteria that is also a natural pathogen of flies. *P. entomophila* has been shown to infect through the oral route resulting in the activation of the fly’s innate immune response and in susceptible flies, resulting in death (Liehl et al., 2006; Siva-Jothy et al., 2018). Similar to the PQ-induced oxidative stress assay, we deployed 3 different feeding regimens: 1)

Pre-stress: flies were fed an AJ-supplemented diet before oral infection; 2) During stress: flies were fed AJ only during infection and 3) Pre & during stress: flies fed AJ before and during infection. The outcome measure was survival 5 days post-infection. Despite their small size, we observed no difference in mortality between control and ELS flies upon infection (Fig. 6). Surprisingly, prophylactic feeding of AJ showed a positive impact on survival upon infection only in ELS flies, with ∼4-fold increase over control flies. Unlike survival under oxidative stress, no further additive gain in resilience was observed when AJ was provided during infection. Although AJ supplementation improves survival of uninfected ELS flies (EU) when provided acutely for 24 hours, it was not statistically significant. The observation that AJ prophylactically benefits only ELS flies may perhaps indicate that when it comes to infection resilience, AJ’s impact might be through epigenetics. Nutritional stress during development is known to reprogram epigenetic signatures (Kubota, 2016). Furthermore, the absence of AJ’s impact on infection survival in control flies as opposed to its benefit during oxidative stress to both control and ELS flies (Fig. 6) suggests that biological mechanisms regulated by AJ to deal with oxidative stress may not overlap with mechanisms that improve innate immunity.

**Figure 6:**
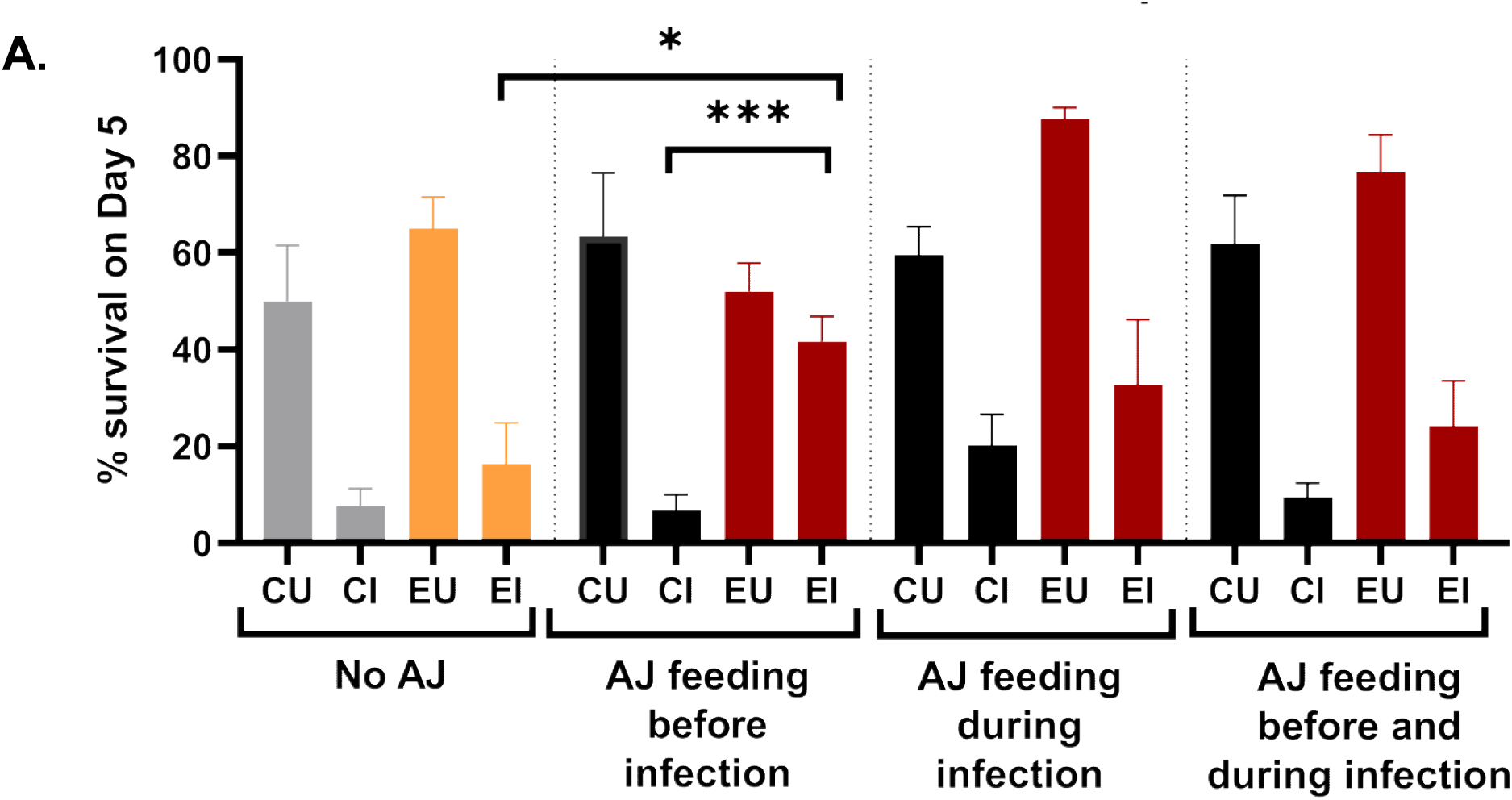
Impact of AJ supplementation on survival upon enteric infection. % of flies alive 5 days post-exposure to P. entomophila. CU: Control Uninfected; CI: Control Infected; EU: ELS Uninfected; EI: ELS Infected n: Control ≥58, ELS ≥ 63. Error bars are ± SEM. * p ≤ 0.05 and *** p ≤ 0.001 calculated using one-way ANOVA. There was no statistical difference between the other bars.

## 4 Conclusion

In this study, we report the effect of dietary Amla juice supplementation on select systemic readouts associated with health in *Drosophila melanogaster* – lifespan, reproduction, resistance to oxidative stress and enteric infection. Outcomes were measured in mated females and males, segregated before assay. Also, supplementation was tested on a fly model for developmental malnutrition, early life starved (ELS) flies. Except for a small but significant negative impact on lifespan of ELS females, chronic Amla supplementation had no impact on control or ELS fly lifespans or development (Fig. 1A,B,D,E). AJ pre-feeding improved egg numbers in ELS flies only (Fig. 1C). Oxidative stress was induced by PQ and three Amla feeding regimens were tested: pre-stress, during stress and a combination of the two (Fig. 2). Regardless of dietary regimen, a large and significant increase in survival to oxidative stress was observed in both control and ELS flies; however, the scale of impact is sex-specific (Fig. 3 and Fig. 4). Upon exposure to *P. entomophila,* a significant impact on survival was only observed in ELS flies subject to Amla pre-feeding (Fig. 6).

## 5 Discussion

Amla is an extremely well-studied fruit as it is associated with several traditional medical claims. What is less appreciated is that Amla is both a food and a medicine.

The latter is more extensively studied, and as detailed in the Introduction, Amla’s mode of action has been studied in cell lines, model systems and human, but primarily in the diseased context. In this study, we attempted to understand if there are any benefits to consuming Amla as a food and specifically, as a nutraceutical to treat adult phenotypes that result from developmental malnutrition. Our data points to specific benefits for prophylactic consumption of Amla. For general wellbeing, Amla appears to confer strong benefits against oxidative stress even when consumed pre-stress. Amla and its constituents have been shown to scavenge oxygen radicals and increase superoxide dismutase activity. Hence, it is expected that when provided during stress Amla will increase survival. The mode of action that underlies it’s prophylactic benefit however is less clear. Furthermore, the positive impact is observed in both sexes. This suggests that the mode of action through which Amla confers resistance to oxidative stress, before the induction of stress, is likely to be fundamental. Here, we only explored expression of superoxide dismutase gene (Fig 5) and found it unchanged. The nrf2 pathway and glutathione recycling are other possible pathways to explore.

In summary, these studies add to the repertoire of knowledge in regard to the benefit of dietary supplementation with Amla (Indian gooseberry). By using an animal model for early-life malnutrition, we demonstrate the potential of adding this ingredient to overcome reproductive deficiencies and improve resilience to oxidative stress, both as a prophylactic and therapeutic. Furthermore, the assays described may be used as a benchmark to test functional food claims associated with the addition of Amla. Although community and clinical trials will be ultimately necessary to understand the efficacy, the absence of toxicity and the advantages of using flies as a screening tool will help to develop new formats and formulations to tackle malnutrition.

## Abbreviations Used

AJ: Amla Juice
ELS: Early Life Starved
ND: Normal Diet
PQ: Paraquat

## Acknowledgements

Support from Rural India Supporting Trust is gratefully acknowledged. We would like to thank Ms. Neha and Mr. Shamsher Singh Mann for donating lab equipment. Thank you Prof Bhushan Patwardhan for supporting this work. Thank you Sadashiv Naik and Vaishali Jadhav for pictures of Amla.

---

**Supplementary Figure 1:**
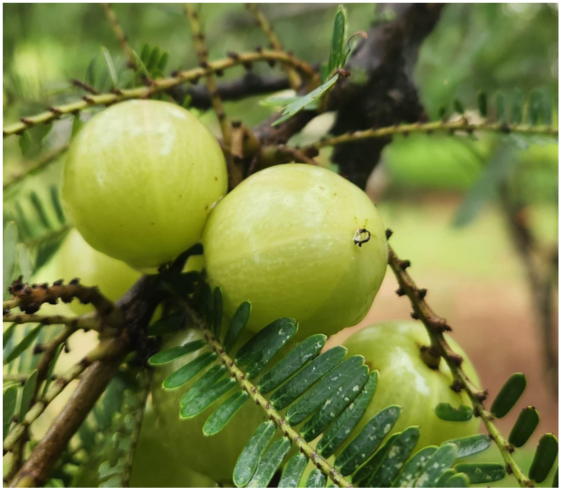
*Phyllanthus emblica L.* (Indian gooseberry, Amla)

**Supplementary Table 1:** Superoxide anion scavenging activity of Amla Juice as measured by its ability to prevent Pyrogallol oxidation, was adapted from Marklund and Marklund (Marklund & Marklund, 1974) and derived equivalence to ascorbic acid is reported.

| Type | % AJ | % Scavenging activity | Equivalence to Ascorbic acid (mmoles) |
| --- | --- | --- | --- |
| Fresh | 1.25 | 50.1 | 0.0032 |
|  | 2.5 | 84.9 | 0.0055 |
|  | 5 | 160.7 | 0.0103 |
| Stored -80°C (3 months old) | 1.25 | 35.8 | 0.0023 |
|  | 2.5 | 88.6 | 0.0057 |
|  | 5 | 162.6 | 0.0105 |

**Supplementary Table 2:**
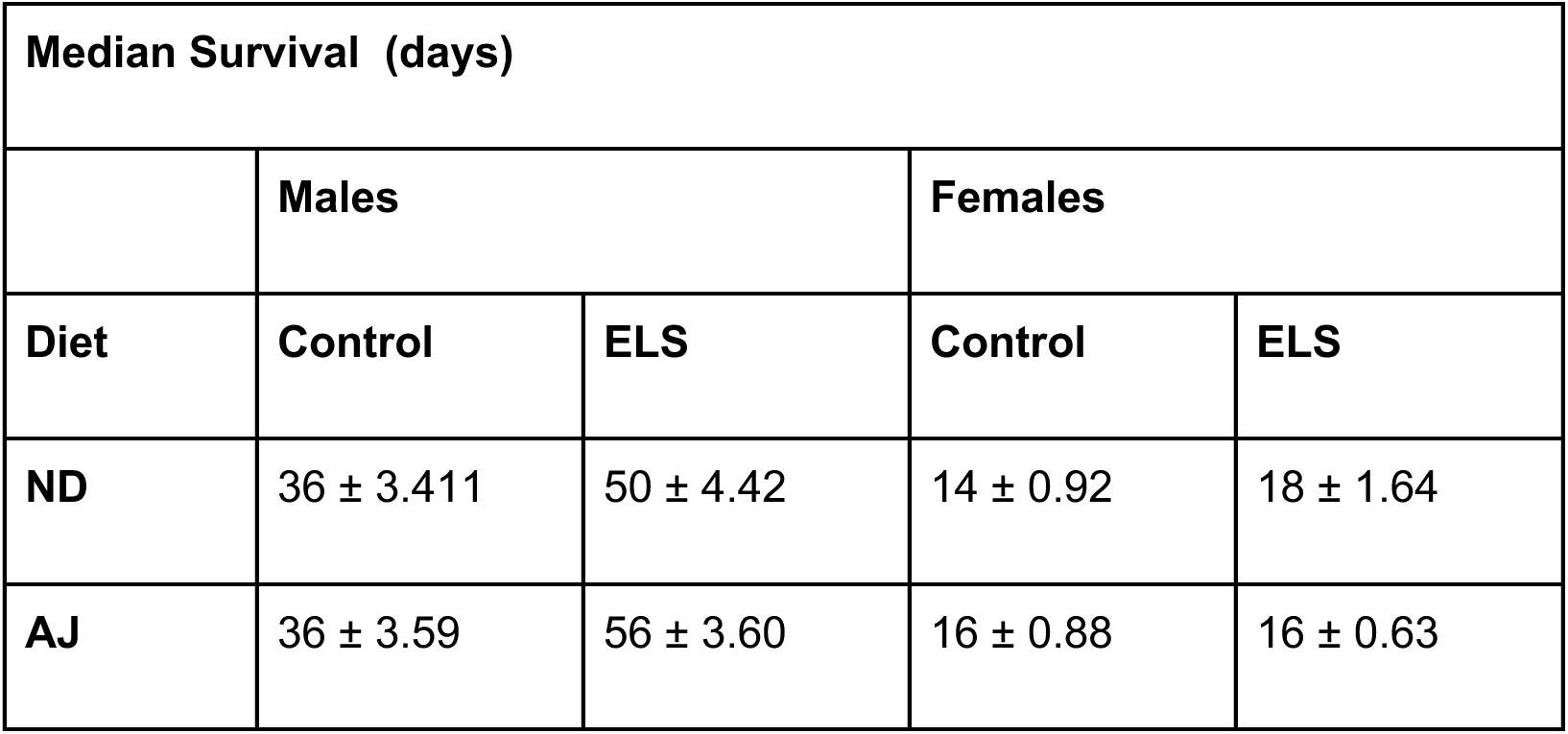
Lifespan of Control and ELS flies: Median survival of flies fed on Amla juice.

| Median Survival (days) |  |  |  |  |
| --- | --- | --- | --- | --- |
|  | Males |  | Females |  |
| Diet | Control | ELS | Control | ELS |
| ND | 36 ± 3.411 | 50 ± 4.42 | 14 ± 0.92 | 18 ± 1.64 |
| AJ | 36 ± 3.59 | 56 ± 3.60 | 16 ± 0.88 | 16 ± 0.63 |

